# Using Repetitive Control to Enhance Force Control During Human-Robot Interaction in Quasi-Periodic Tasks

**DOI:** 10.1101/2021.05.09.443322

**Authors:** Robert L. McGrath, Fabrizio Sergi

## Abstract

We investigated the use of repetitive control (RC) to enhance force control during human-robot interaction in quasi-periodic tasks. We first developed a two-mass spring damper model and formulated three different RCs under force control: a 1^st^ order RC (RC-1), a 3^rd^ order RC designed for random period error, and a 3^rd^ order RC designed for constant period error. Then, we quantified the performance of these three RCs through simulations and experiments conducted on a bench top linear platform, subject to nominal cyclical inputs (input signal and fundamental frequency: 0.5 Hz), and subject to inputs with random and constant period errors. Moreover, we compared the performance achieved with the RCs with those achievable with a passive proportional controller (PPC), subject to known theoretical limits for passivity and coupled stability.

In both simulated and real-world experiments, the root mean square force error under nominal conditions was reduced most effectively by the RC-1 to 0.7% and 12.9%, respectively, of the error achieved with the PPC. Subject to inputs with constant period errors, RCs performed better than PPC for period error values below 0.05 Hz, with the RC-1 performing significantly better than both 3^rd^ order RCs. Subject to inputs with random period errors, all RCs performed better than PPC up to 0.11 Hz of frequency error. Our results indicate that RC can successfully integrated in force control schemes to improve performance beyond the one achievable with a PPC, in the range of period variability expected in applications such as walking assistance and rehabilitation.

## I. INTRODUCTION

Transparency, or the achievement of low interaction forces during unassisted movements, is a necessary quality of effective human-interacting robots as low-interaction forces will allow for natural movement patterns [1], [2]. Force feed-back is a simple method implemented on previous robotic platforms [3], [4], but has limitations due to the fact that the input of the plant does not change until error is already observed. Furthermore, stability of force-feedback system crucially depends on an accurate knowledge of the dynamics of the controlled system, knowledge that can be difficult to obtain in practice. In fact, a system using high control gains tuned to achieve high transparency performance may not be passive, and thus could be unstable during interaction with a percentage of the user population. Passivity is a desirable condition for the dynamics of a human-interacting robot, as it guarantees the stability of the robot when interacting with a range of environmental conditions [5], [6]. The trade-off between stability and performance is particularly an issue on robotic platforms with high reflected inertia and therefore high interaction forces [3]. As such, there is a need for novel controllers capable of reducing interaction forces while remaining stable for a robust range of conditions such as those encountered during human-robot interaction.

Many scenarios involving physical human-robot interactions, such as robot-assisted manufacturing or robotic rehabilitation, are centered on tasks that are nearly cyclical such as repetitive point-to-point movements or walking. Specifically, the walking gait has been identified as a quasi-periodic task, in which the motion is continuous, approximately periodic, but with a non-zero variance in period duration [7]–[10]. The quasi-periodic nature of these tasks could be exploited for the purposes of enhancing the performance of controllers supporting humans during these tasks via force feedback. In fact, if the input provided by the subject at the current cycle of operation will be nearly equal to the input at the next cycle, a controller could be designed to compensate for future force error.

In previous Bowden cable driven exoskeleton work, iterative learning control (ILC) [11] has been utilized to learn quasi-periodic cable tensioning patterns for the gait cycle [12]. The implementation of this learning controller was possible due to the cable slacking in the swing phase of the gait cycle, creating a reset point between periods. However, not all human-interacting robots have this possibility, and in most cases the interaction forces are continuous signals with the same periodicity as the resulting motion. The repetitive controller (RC), intended for periodic and continuous operation in the time domain, could be a feasible option for human-interacting robots such as gait training exoskeletons during continuous operation [13], [14]. RC is a feedforward control method for reducing error of the current task cycle based on the error observed in the previous task cycles, and is suitable for continuous operations. In robotics, while ILC have also been used in force control [11], [12], [15], RCs have been primarily applied to trajectory control [16], [17]. However, we are not aware of any applications of RC in force control.

RCs require some knowledge of the dynamics of the system they are applied to, and have traditionally been applied to systems with highly regular periodicity [18]. In human-interacting robots, the dynamics of the coupled system may not be known with sufficient accuracy due to between-subject variability [19]. Also human tasks such as walking gait are quasi-periodic, in which stride-by-stride variation in gait spatiotemporal parameters is on the order of 3-5% [20]. There are formulations of RCs (high-order controllers) which combine information measured on multiple previous cycles to improve robustness to input period error [21]–[24], but it is unclear how these formulations would respond to variability in input parameter period. Also, to the best of our knowledge, RCs had not been used in force controllers for human-robot interaction, which are known to present challenges inherent with the coupled stability of the system [5], [6]. The novel contribution of this work is the application of RC and high order RC to force control in a human-interacting robot, and analysis of the performance of the controller in the context of quasi-periodic tasks.

In this work, we first present formulations for RCs that use force feedback to reduce the interaction forces of a human-interacting robot. We test the performance of these RCs via simulations applied to a two-mass spring damper system classically used to study controllers for human-interacting robots under force control. We then simulate behavior in a case where we assume perfect knowledge of system parameters and perfect periodicity of subject inputs. Then, we conduct simulations in non-ideal conditions to assess the robustness of different formulations of the controller to variations in parameters of the system dynamics, and to variations in the period of the inputs that would be applied by a subject. Moreover, we validate our simulations via experiments conducted on a bench-top linear actuator platform designed for force feedback human-interaction. Experiments are conducted in nominal conditions, to assess stability and reduction in interaction force, as well as under constant and random signal period error matching conditions tested in simulation.

## II. METHODS

### A. Mechanical System for Simulation

We used a two-mass model classically studied in force control for human-interacting robots [5], [6] to evaluate the behavior of the RCs under force control. The mechanical system considered in this paper for simulation, shown in Fig. 1, consists of two masses connected by a spring and damper in parallel. In the general case of a physical human-robot interaction, the proximal *M*_1_ and distal *M*_2_ masses are representative of the motor rotor and end-effector, respectively. The spring *K* and damper *B* in parallel are representative of the transmission dynamics. The distal mass interacts with an environment modeled as a velocity source *V*_2_, as might theoretically be imposed by a human participant, with resulting environmental force *F*_*E*_. The proximal mass has an applied control action motor force *F*_*A*_ as specified by controller architecture explained in the following sections.

**Fig. 1:**
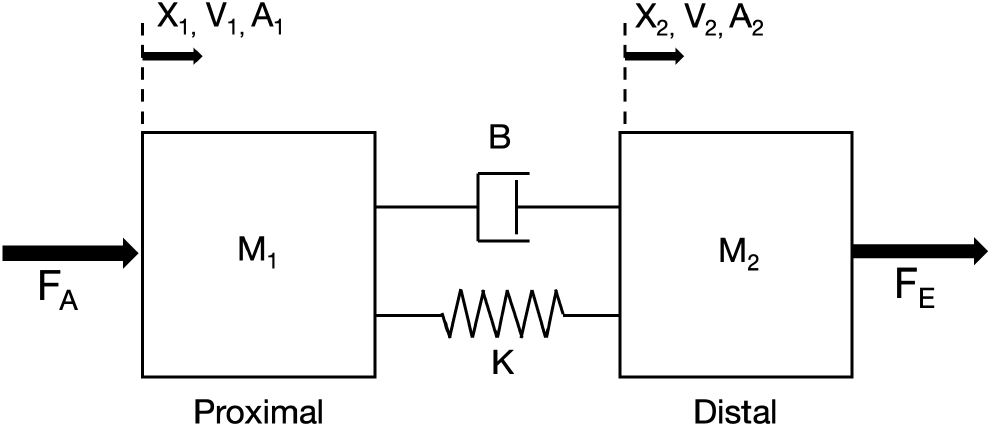
Mechanical system of interest.

**Fig. 2:**
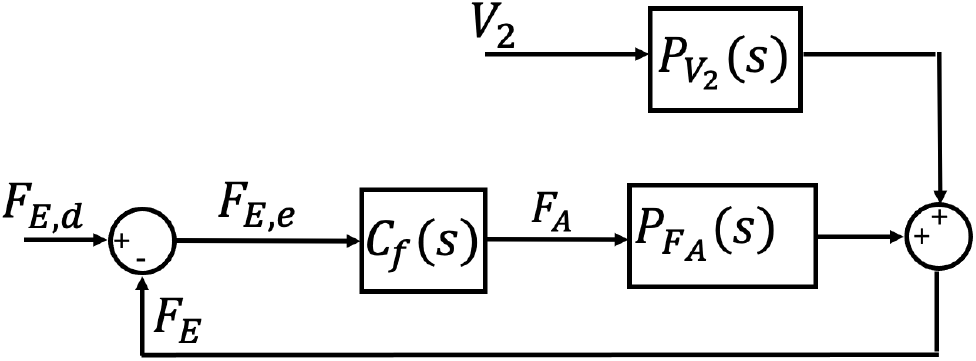
Schematic of a standard zero-force controller with inner force-feedback loop.

### B. Zero-Force Controller

This controller is designed with a desired environmental force *F*_*E,d*_ of zero while subject to a distal velocity disturbance *V*_2_ (Fig. 1). The equation relating the environmental force *F*_*E*_ to the distal velocity *V*_2_ and control action motor force *F*_*A*_ and plant transfer functions is given as:

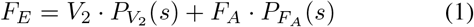

where the plant transfer functions 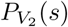 and 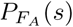 take the form:

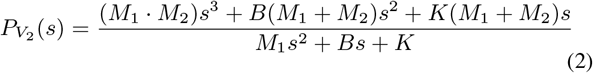

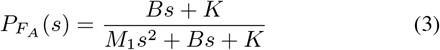

The control action motor force *F*_*A*_ applied to the proximal mass is obtained by taking the error force *F*_*E,e*_ between the measured environmental force *F*_*E*_ and the desired environmental force *F*_*E,d*_ and passing it through the control block transfer function *C*_*f*_ (*s*). We are requiring the controller to meet the condition of passivity, to ensure stability when interacting with a range of environmental conditions [6]. As such, the control block transfer function *C*_*f*_ (*s*) is required to satisfy the passivity requirement of:

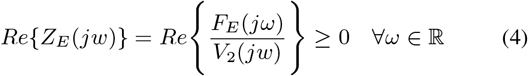

This leads to the following expression for the passive upper limit on the proportional gain *K*_*P*_ :

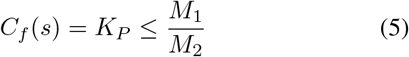

Although the expressions for the transfer functions are derived in the s-domain, the transfer functions are simulated in the z-domain in the following sections using a zero order hold (*T*_*s*_ = 0.001*s*)

### C. Repetitive Force Control

In an RC, the error of the previous cycle is applied through a feed-forward system to compensate for the anticipated error of the current cycle. The general form of the first order RC [25] is as follows:

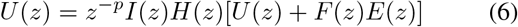

The RC is designed and implemented in the *z* domain. The controller’s output *U* (*z*) and error *E*(*z*), operated on by a compensator *F* (*z*), from the previous cycle (given delay *z*^*−p*^) are combined to produce the output at the current cycle. The interpolator *I*(*z*) is necessary if the number of time steps in a cycle *p* is not an integer and the zero phase low-pass filter *H*(*z*) can be applied if a frequency cutoff of the learning process is necessary.

The transfer function form of the general 1^st^ order RC (RC-1), converting the error input *E*(*z*) to control output *U* (*z*), is:

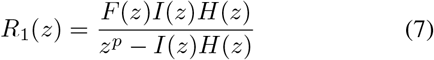

Given the scope of this work, the interpolator *I*(*z*) has been set to unity, as the capability of specifying a nominal frequency that required a fractional number of points in unnecessary. Also, we have added a learning gain parameter *ϕ* to provide weight to the compensatory action which is constrained such that:

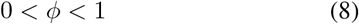

This yields the following transfer function for an RC-1:

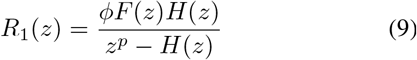

The closed loop transfer function *G*(*z*) is defined as:

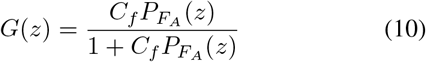

We also implemented an additional controller, a 3^rd^ order RC, with generic transfer function form:

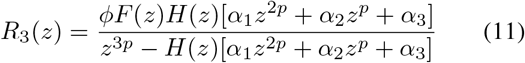

The parameters *α*_1,2,3_ provide weights to the compensatory action resulting from the previous three cycles and is constrained such that:

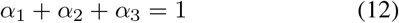

For this work, we have implemented plug-in type RCs [26], [27], as shown in Fig. 3, where the RC transfer function is located in parallel to a standard feedback loop.

**Fig. 3:**
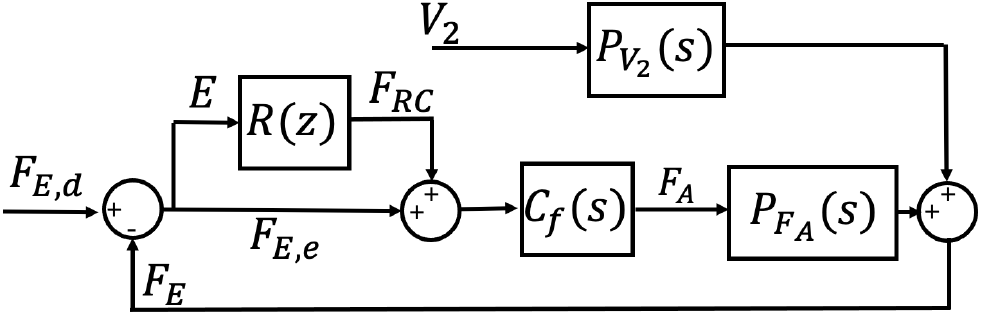
Schematic of the zero-force controller with applied plug-in type repetitive control.

We opted for this form of controller, as opposed to the cascade-type RC, because the plug-in type could be conveniently enabled or disabled without disruptive transient of force error. This capability was deemed to be essential for safety-critical applications such as human-robot interaction.

### D. Repetitive Controller Tuning

For these RCs, the compensator *F* (*z*), a finite impulse response digital filter, is specified by two design terms *m* & *n*, where the compensator has *m* zeros and *n − m* poles. We know that the closed-loop system with added RC (plug-in or cascade) and zero phase low-pass filter is stable if the following expression holds:

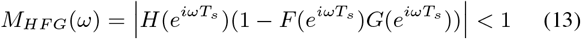

To determine a suitable compensator, the following expression is minimized [27] over the frequency range of interest utilizing *fmincon* in MATLAB (MathWorks Inc, Natick, MA):

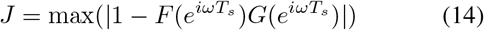

Two sets of 3^rd^ order RC weight parameters were based on previous work [28] and tuned empirically for performance. These sets of parameters *α*_1,2,3_ were designed specifically for accommodating random period error (RPE) and constant period error (CPE) and assigned as [3/6, 2/6, 1/6] and [1, -1, 1], respectively. The learning gains *ϕ* for the 3^rd^ order RC designed for RPE (RC-RPE) and 3^rd^ order RC designed for CPE (RC-CPE) controllers were determined empirically and set as 0.50 and 0.25, respectively.

### E. Nominal Simulation Evaluation

All controller simulations were implemented via Simulink models with a distal velocity signal obtained as the time integral of a unit magnitude acceleration signal, sampled at a specified interval (*T*_*s*_ = 0.001s). Each nominal simulation began with 10 seconds (5 cycles) of passive proportional control (PPC) followed by the plug-in type RC engaged for 60 seconds (30 cycles). We used the outcome measure of root mean squared value of force error 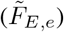, quantified for each cycle of the source signal.

### F. Simulation Robustness Analyses

#### 1) Model Parameter Error

The compensator optimization process, based on a model of the plant transfer function 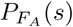 and the passive control block *C*_*f*_ (*s*), determines the stability of the RCs. However, in practical implementation of the RCs, there will exist error in the plant transfer function estimate relative to the actual physical plant. As such, it was important to asses the stability of the RC when there was error present in the modeled transfer function. In order to assess the robustness of the RC to model error, we assessed the convergence of the controllers when the compensator was designed with an ideal plant model and was applied to plant models with various magnitude of error applied to the individual parameters (i.e., *B, K, M*_1_, *M*_2_).

For each of the three RCs, we ran one hundred simulation repetitions for ten values of parameter error (i.e., 0%, 10%, …, 90%). As such, if a nominal value was 9 kg and the parameter error was equal to 10%, the value simulated in each of the 100 repetitions was randomly sampled from a uniform distribution with margins [8.10, 9.90] kg. Each simulation started with five cycles of PPC followed by 245 cycles of plug-in type RC. The convergence of each simulation was assessed by the relative magnitudes of the 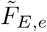 of early RC (mean of cycles 6-11) and late RC (mean of the last 10 cycles). If the late 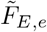 values was less than the early 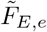 value, the simulation was considered to converge.

#### 2) Source Signal Random Period Errors

The controller is subject to the imposed distal velocity source, modeled as a sinusoidal waveform. Quasi-periodic tasks are characterized by cycle-by-cycle variability in cycle period. As such, there will be an error between the true period duration of the quasi-periodic source and the period assumed by the controller. This error can be randomly distributed across cycles (random period error, RPE).

To assess the robustness of the RCs subject to RPE, we assessed the 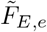 of the controllers when the period of the distal velocity signal varied randomly with a uniform distribution. For each simulation of the RPE analysis, the frequency of each individual cycle of the applied sinusoid was drawn from a uniform distribution of with a mean of 0.5 Hz and variable half-width. The half-width was changed to simulate different levels of RPE and ranged between 0 Hz to 0.45 Hz (in increments of 0.005 Hz). Each simulation began with 10 cycles of PPC followed by 190 cycles of plug-in type RC. The 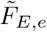 of each controller was assessed: for PPC (mean of cycles 2-10) and for the RCs (mean of last 100 cycles).

#### 3) Source Signal Constant Period Errors

The error between the true period duration of the quasi-periodic source and period of the assumed controller can also be constant (constant period error, CPE) in the case of period drift. In order to assess the robustness of the RCs to CPE, we assessed the performance of the controllers when the cycle period was a consistent non-nominal value for the duration of the simulation. This is a worst-case scenario to assess the robustness of an RC to a discrepancy between the perturbation period and the period assumed by the RC. We assessed 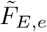 at a range of 51 different distal velocity frequencies (i.e., 0.25 - 0.75 Hz), with a constant spacing of 0.01 Hz. Each simulation, at a specified distal velocity frequency, began with 5 cycles of PPC followed by 45 cycles of plug-in type RC. The 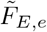 of each controller was assessed: for PPC (mean of cycles 2-5) and for the RCs (mean of last 5 cycles).

### G. Experimental Platform

Force feedback control was implemented on a linear actuation platform shown in Fig. 4. The platform consists of a PRM style 8mm diameter x 8mm lead ballscrew (Thomson Industries, Inc., USA) driven by a belt and pulley system (1:5 gear ratio) and EC 4-pole 30 motor (Maxon Motor; Sachseln, Switzerland) The nut of the ballscrew is attached to the carriage of a 500 series linear guide (Thomson Industries, Inc, USA) by custom 3D printed components to form an assembly. The assembly includes a Mini27Ti 6-axis transducer (ATI Industrial Automation, USA) attached to a handle such that the human-robot interaction force parallel to the direction of linear actuation can be measured. The transducer signals are acquired with a PCIe-6321 DAQ Card (National Instruments). The motor is controlled by an EPOS2 50/5 motor controller (Maxon Motor; Sachseln, Switzerland) with accompanying motor control software suite (EPOS2 Studio) which interfaces with a Q8 DAQ board (Quanser, Canada) for current command and encoder reading acquisition. The hardware is controlled by custom control software coded in Simulink & MATLAB (Mathworks, Inc, USA) with Quarc realtime control blocks (Quanser, Canada) running at a sample and update rate of 1k Hz. As seen in Fig. 4, behind the robotic platform we positioned a computer monitor displaying a visual metronome. This was used as a visual cue for the operator to impose controlled roughly sinusoidal waveforms of distal position, and therefore velocity. In all experiments, the operator received visual feedback about the desired position and tried their best to apply the force required to achieve the desired kinematics.

**Fig. 4:**
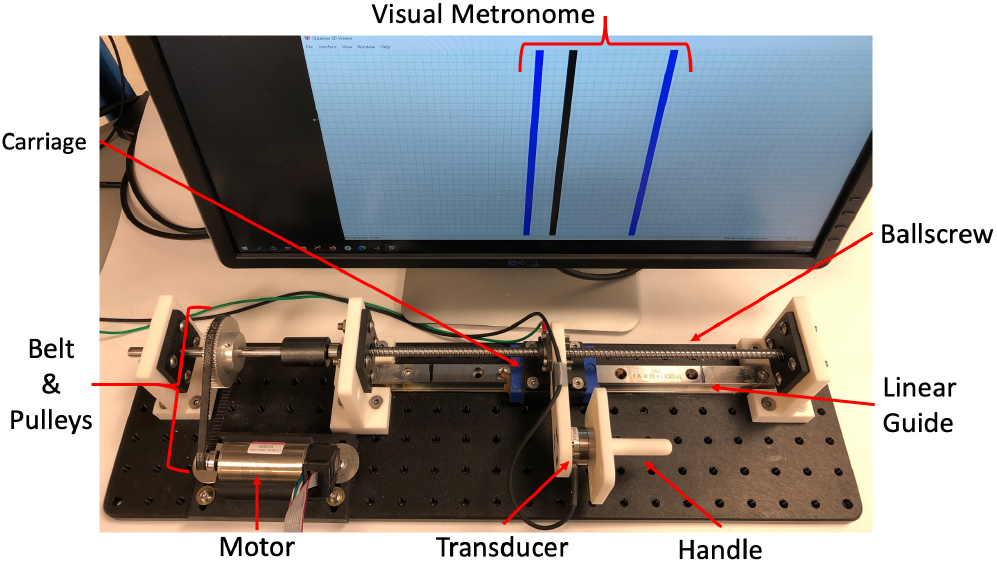
Experimental setup consisting of a pulley driven linear actuator with sensorized handle for human-robot interaction.

### H. Tuning and Characterization for Validation

A proportional force feedback controller was tuned for the highest gain enabling the stability of the robotic platform with the worst case environmental condition for stability. For characterization, the experimental platform was commanded a Schroeder Multisine force signal with the robot in closed-loop proportional feedback control while the end effector was physically blocked. The commanded Schroeder Multisine and resulting measured force were processed utilizing the *tfestimate()* function in MATLAB to produce the magnitude and phase numerical estimates of the closed loop transfer function *G*(*z*) over the characterizable frequency range. With this process, a numerical estimate of *G*(*z*) over a limited frequency range is obtained. The estimated *G*(*z*) is then used to design the compensator F(z) based on eq. (14). The resulting RC design process only differs from the previously described methods in sections II-C & II-D in two aspects. First, the compensator is specifically optimized for performance over the characterized range of frequencies [18] and therefore might perform poorly at frequencies outside this range, specifically higher frequencies. Second, to address this, the digital zero-phase low pass filter *H*(*z*) is designed to cutoff the learning of the designed RCs above the upper end of the characterized range of the closed loop transfer function *G*(*z*).

### I. Nominal Performance Validation

Each simulation began with 20 seconds of PPC followed by plug-in type RC engaged for 90 seconds. As in simulations, the primary outcome measure was the 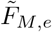 for each cycle according to the velocity prescribed by the visual metronome.

### J. Robustness Validation

#### 1) Applied Random Period Error

For each trial of the applied RPE analysis, the frequency of each individual cycle of the operator applied sinusoid under visual feedback was drawn from a uniform distribution with a mean of 0.5 Hz and variable half-width. The half-width was changed for each trial and ranged between 0 Hz and 0.125Hz (in increments of 0.025 Hz). Each trial began with 20 seconds for calibration and startup, 20 seconds for simple proportional control, and 90 seconds of plug-in type RC. The 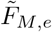 was assessed as a percentage: the mean of the 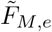 of the last 30 cycles of RC divided by the mean of the 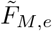 of the 10 cycles of PPC. The values of 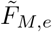 were plotted versus the calculated half-width of the trial: an estimate of the continuous uniform distribution, based on the variance of *V*_2_, derived from model calculations utilizing the measured *F*_*M*_, *V*_1_, and model parameters (i.e., *B, K*, and *M*_2_). A general linear model was fit to the data with the RC type and half-width as fixed effects. Pairwise comparisons were performed between the modeled controller values at a low, medium and high half-width value.

#### 2) Applied Constant Period Error

We assessed 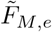 at a range of 5 different distal velocity frequencies (i.e., 0.50 -0.60 Hz), imposed by an operator under visual feedback. Each simulation, at a specified distal velocity frequency, began with 10 seconds of PPC followed by 90 seconds of plug-in type RC. The 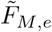 of each controller was assessed as a percentage: the mean 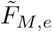 of the last 30 cycles of RC divided by the mean 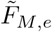 of the 10 cycles of PPC. The values of 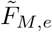 were plotted versus the calculated frequency of *V*_2_ derived from model calculations utilizing the measured *F*_*M*_, *V*_1_, and model parameters (i.e., *B, K*, and *M*_2_). A general linear model was fit to the data with the RC type and frequency error (difference from nominal frequency) as fixed effects. The model also included the interaction term between the fixed effects, and an explanatory variable for frequency error squared. Pairwise comparisons were performed between the modeled controller values at a low, medium and high frequency error values.

## III. RESULTS

### A. Repetitive Controller Tuning for Simulation

Prior to performing simulations, the compensator was designed according to the specified default set of parameters given in Table I. This set of parameter were derived from parameterizing the experimental platform as a two-mass spring-damper system. The two mass values were based on the lumped equivalent masses per CAD model calculations, with the pulley as the point of separation of the two masses, in the linear domain of the end-effector. The spring constant was estimated based on the frequency of vibration of the proximal mass during an applied step input and blocked configuration of the distal mass (Fig. S1). The damping coefficient was determined through estimation of the damping ratio based on inspection of the step input transient response.

**TABLE I:**
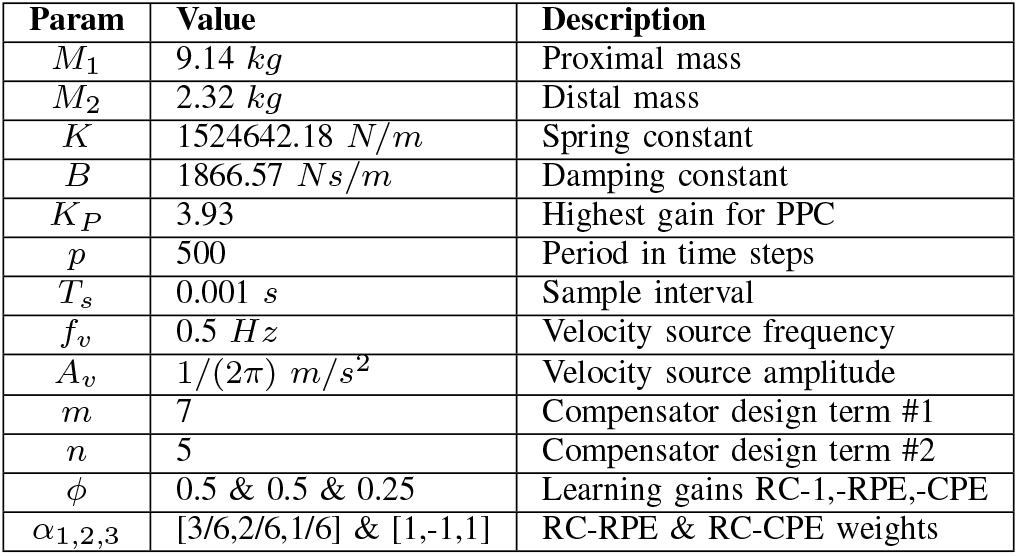
The default set of simulation parameters

In Fig. 5, the optimized compensator is shown and the resulting asymptotic stability obtained when coupling the compensator with the nominal plant for both the low pass filtered and unfiltered systems (*M*_*HFG*_ *<* 1 and |1 − *FG* |*<* 1) for all frequencies of interest.

**Fig. 5:**
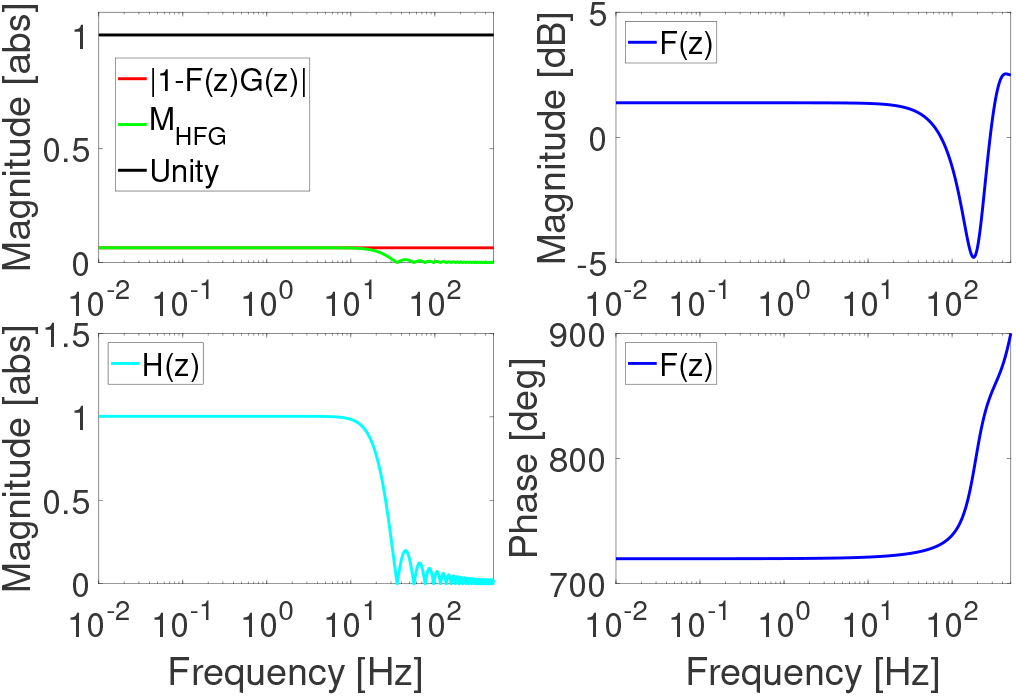
(Top left) Asymptotic stability of the compensator when coupled with the nominal plant and zero-phase low pass filter, (bottom left) the magnitude of the low pass filter, and (top right) the magnitude and (bottom right) phase of the compensator.

### B. Nominal Simulation Performance

The nominal simulations for plug-in type RC-1, RC-CPE, and RC-RPE applied to the zero-force controller are shown in Figs. 6-8, respectively. After the PPC quickly achieves steady state, the RCs are applied and achieve a new steady state after several cycles of learning. The RC-1 reduces the 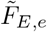 to below 5% of the 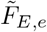 achieved with PPC after 6 cycles of learning, while the RC-CPE and RC-RPE require 21 and 10 cycles, respectively, to achieve the same 95% reduction of 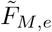.

**Fig. 6:**
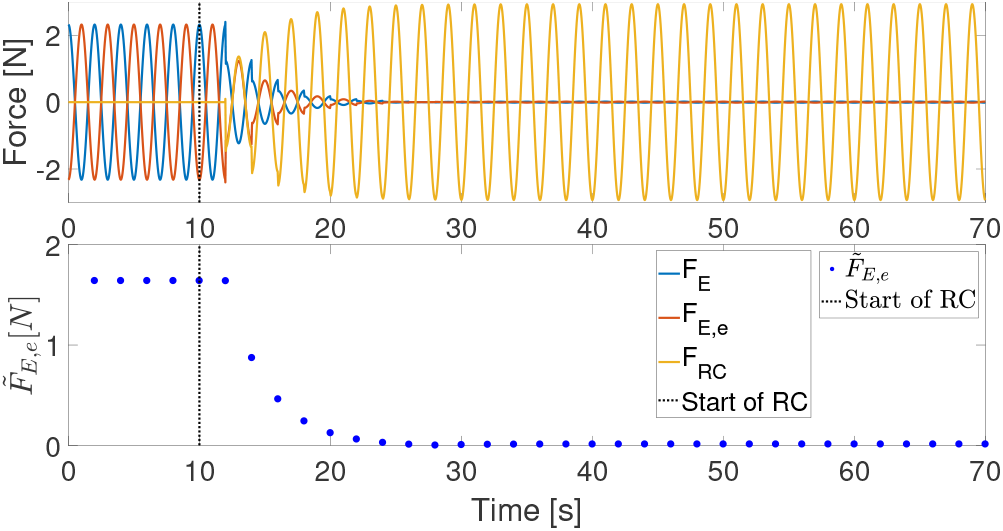
(Top) simulation of the plug-in type 1^st^ order repetitive controller applied after 10 seconds of passive proportional control. (Bottom) the RMS force error for each cycle.

**Fig. 7:**
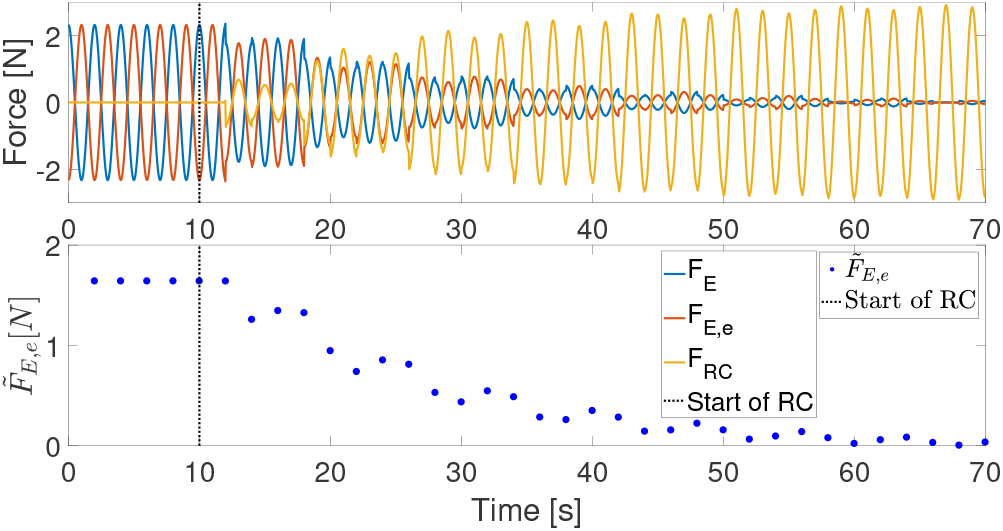
(Top) simulation of the plug-in type 3^rd^ order repetitive controller, designed for constant period error, applied after 10 seconds of passive proportional control. (Bottom) the RMS force error for each cycle.

**Fig. 8:**
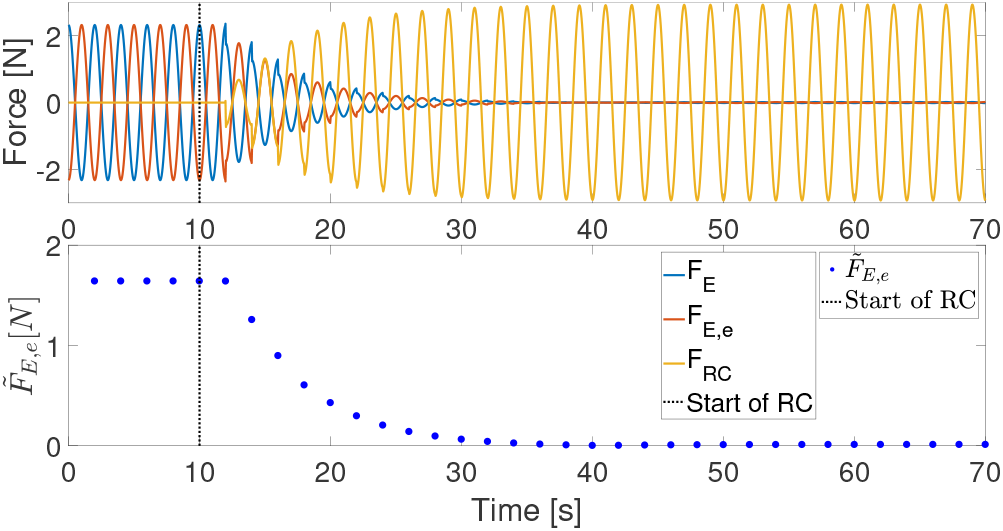
(Top) simulation of the plug-in type 3^rd^ order repetitive controller, designed for random period error, applied after 10 seconds of passive proportional control. (Bottom) the RMS force error for each cycle.

### C. Simulation Robustness Analyses

#### 1) Model Parameter Error

As shown in Fig. 9, all controllers performed with 100% of simulations converging at nominal parameter values (0% error) and maintained 100% convergence through 40% parameter error. All three controllers performed similarly at higher percentages of parameter error in which performance convergence dropped to 60-80% by 90% plant parameter error.

**Fig. 9:**
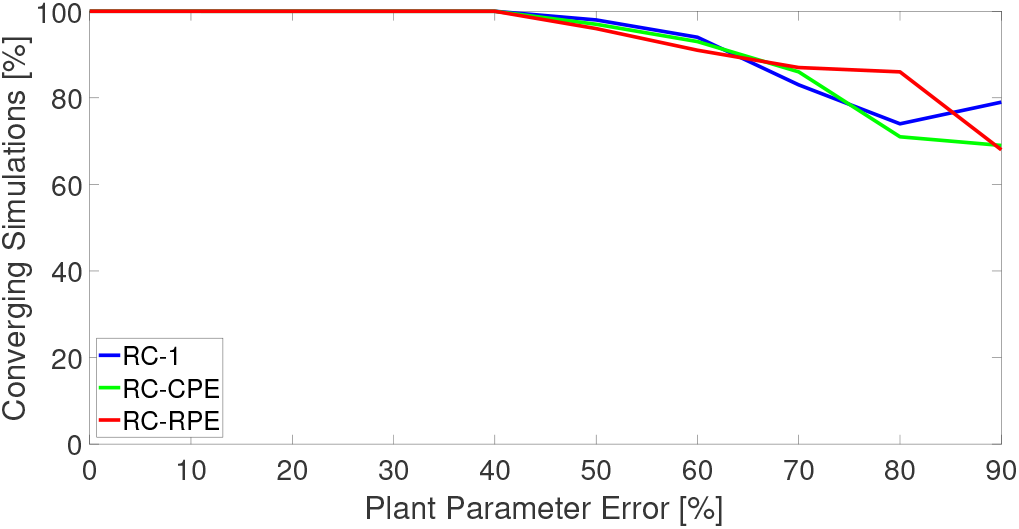
Rate of convergence of simulations under conditions of parameter error for the three repetitive controllers.

#### 2) Source Signal Random Period Error

Under RPE conditions, the PPC performed uniformly across (Fig. 10) all velocity RPE conditions. All RCs performed better than the PPC controller for half-width values below 0.15 Hz (or 30% of the source input frequency). In this range of low RPE values, the RC-1 performed better than the RC-CPE & RC-RPE. At higher half-width values (i.e., above 0.20 Hz or 40% of the source input frequency), instead, the RCs performed worse than the PPC. In this range of high RPE values, the RC-RPE performed better than the other two RC conditions.

**Fig. 10:**
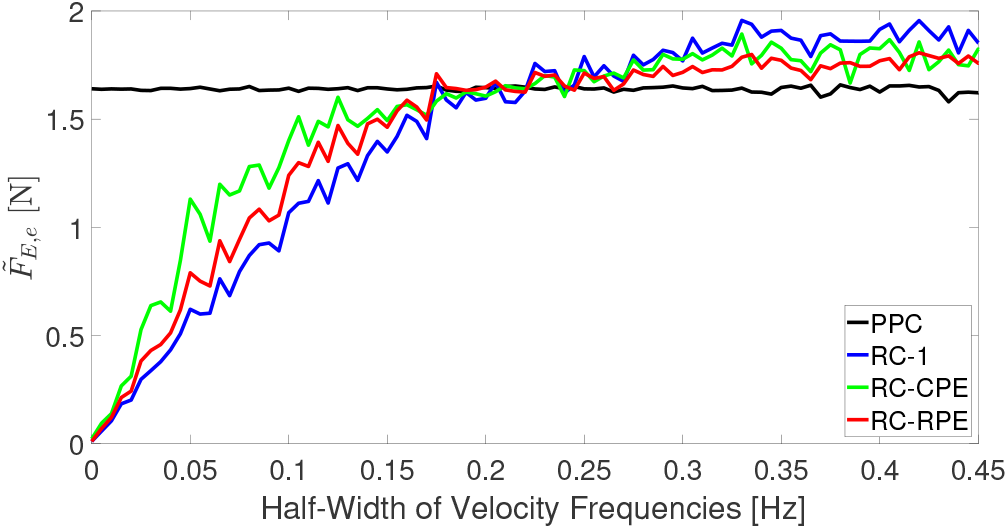
RMS force error for the four force controllers under random error of the period of the distal velocity signal.

#### 3) Source Signal Constant Period Error

The RC-1 performed the best at frequency values near nominal (*±* 0.05Hz, or 10% of the source signal period) and the RC-CPE performed the best at mid-range frequency values (*±* 0.05 Hz to 0.125 Hz). The RC-RPE performed better than the other two RCs only at frequencies far from nominal (*±* 0.125 Hz or 25% of the source signal period), Fig. 11. However, all three RC conditions perform worse than PPC at these far from nominal frequencies.

**Fig. 11:**
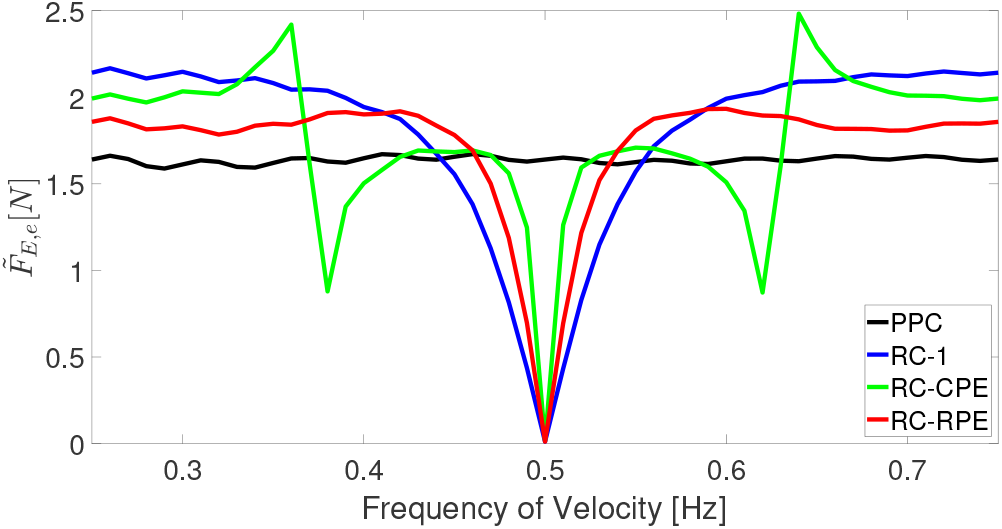
RMS force error for the four force controllers under constant period error of the distal velocity signal.

### D. Tuning and Characterization for Implementation

The *K*_*P*_ gain established for stable control during experiments was 1; a value determined from the worst case environmental condition of coupled stability with a blocked end-effector. The Schroeder Multisine force command signal ranged in frequency content from 0.005Hz to 3 Hz. The resulting estimate of *G*(*z*) had high coherence (*>*0.95) in the range of [0.05 3] Hz (Fig. S2). Given an *m* of 7 and *n* of 5, we achieved an asymptotically stable compensator over the characterized frequency range as can be seen in Fig. S3.

### E. Nominal Performance Validation

The RC-1 required 3 cycles to drop below 20% 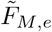, reached a minimum of 8.94%, and maintained a mean of 12.03% in the final 30 cycles (Fig. 12). The RC-RPE required 5 cycles to drop below 20% 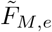, reached a minimum of 8.61%, and maintained a mean of 12.84% in the final 30 cycles (Fig. 13). The RC-CPE required 10 cycles to drop below 20% 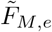, reached a minimum of 10.60%, and maintained a mean of 15.25% in the final 30 cycles (Fig. 14).

**Fig. 12:**
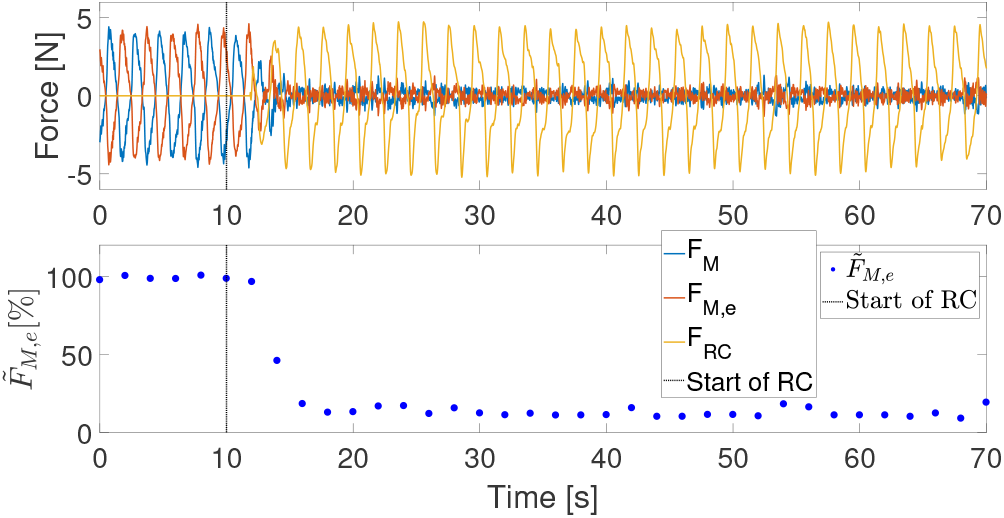
1^st^ order repetitive controller under nominal hand applied perturbations.

**Fig. 13:**
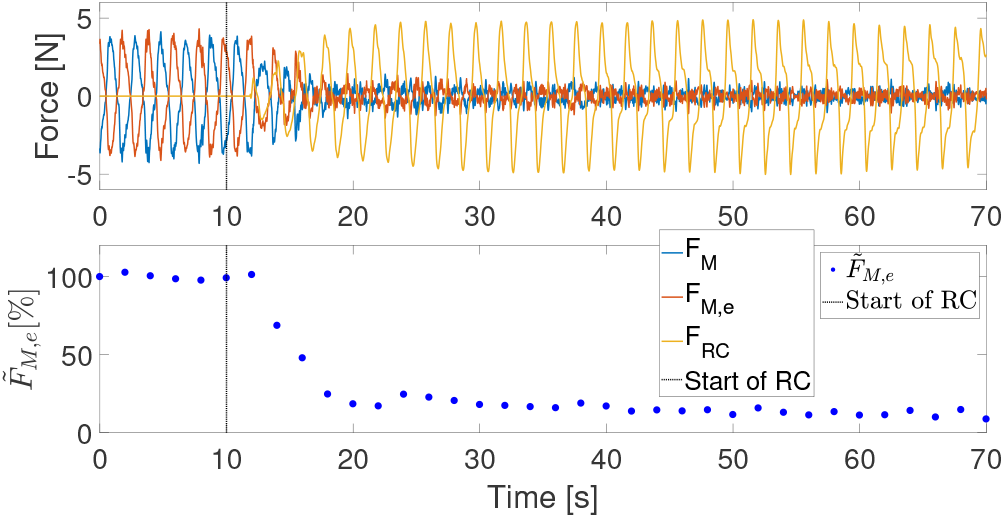
3^rd^ order repetitive controller designed for random period error under nominal hand applied perturbations.

**Fig. 14:**
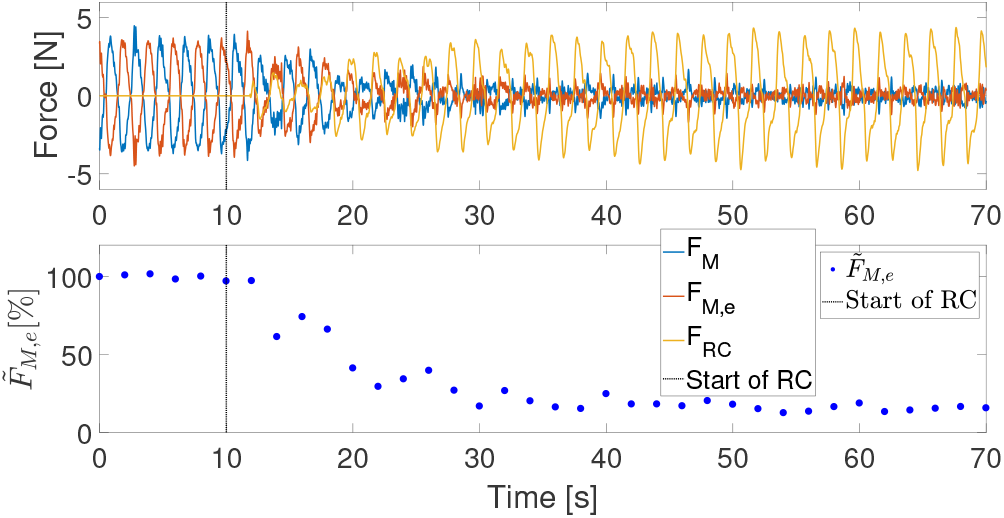
3^rd^ order repetitive controller designed for constant period error under nominal hand applied perturbations.

### F. Robustness Validation

#### 1) Applied Random Period Error

Based on the 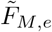 data collected at different source velocity frequency variance values (see Fig. 15), the RC-1 performed generally better than the RC-CPE & RC-RPE, but all three RCs performed equal to or better than PPC in the frequency range of evaluation. According to the post-hoc comparisons resulting from the general linear model fit, at the evaluation half-width of 0.04 Hz, there were no significant difference between controllers (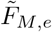: RC-1: 17.45*±* 4.60%, RC-CPE: 20.27 *±* 4.97%, RC-RPE: 21.35 *±* 4.83%). At the evaluation half-widths of 0.08 Hz and 0.11 Hz, the RC-1 (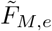: 46.41 *±* 3.51% & 68.14 *±* 5.54%) and the RC-RPE (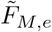: 52.02 *±* 3.55% & 75.03 *±* 5.90%) performed significantly better than the RC-CPE (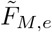: 62.32 *±* 3.50% & 93.86 *±* 5.79%) for all four comparisons (p*<*0.05). At the half-widths of 0.08 Hz and 0.10 Hz, there were no significant differences in performance between the RC-1 and the RC-RPE.

**Fig. 15:**
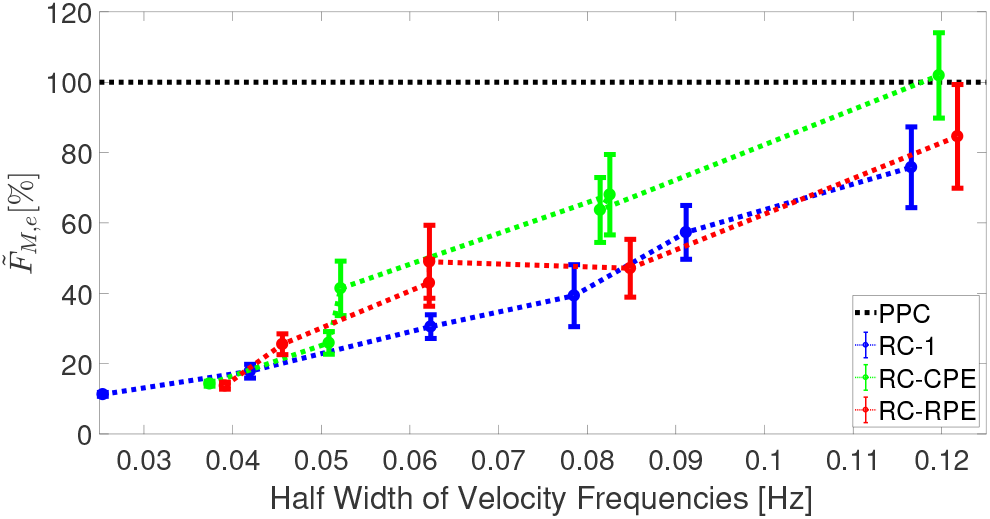
RMS force error as a percentage of passive proportional control for the three repetitive controllers under random error of the period for hand applied perturbations.

#### 2) Applied Constant Period Error

Under conditions of constant period error (Fig. 16), the RC-1 and RC-RPE performed better than RC-CPE at frequency error less than 0.05 Hz. At 0.05 Hz, the PPC performed equal to or better than all three RCs. Above 0.05 Hz, the RC-CPE had similar performance as PPC, and performed the best of all three RCs.

**Fig. 16:**
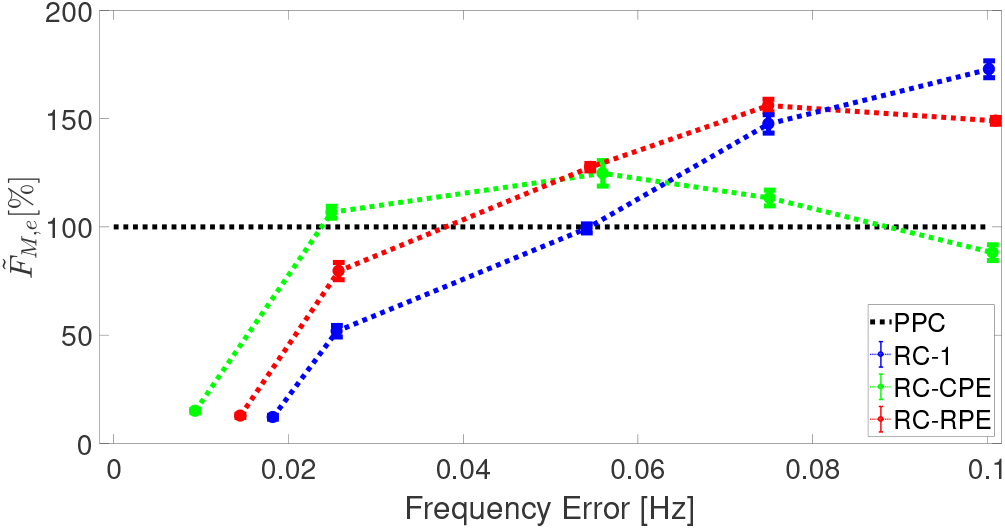
RMS force error as a percentage of passive proportional control for the three repetitive controllers under constant error of the period for hand applied perturbations.

According to the model, at the evaluation frequency error of 0.02 Hz, the 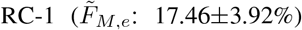 performed significantly better (p*<*0.05) than the 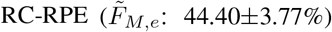, and both performed significantly better (p*<*0.05) than the 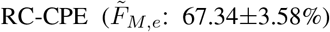. At the evaluation frequency error of 0.06 Hz, the 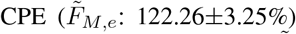 performed significantly better (p*<*0.05) than the 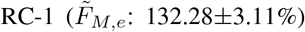, and both performed significantly better (p*<*0.05) than the 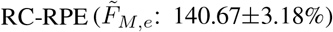. At the evaluation frequency error of 0.10 Hz, the 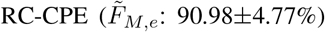 performed significantly better (p*<*0.05) than the 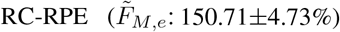, and both performed better (p*<*0.05) than the 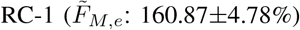.

## IV. DISCUSSION AND CONCLUSIONS

In this paper, we have shown that repetitive force control can be successfully applied to a passive zero-force controller for the reduction of interaction force error 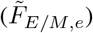 in both simulation and implementation. We have established that repetitive control (RC) is robust to plant parameter error in simulation and sufficiently robust in implementation to achieve convergence. Furthermore, we have quantified the effects of constant and random period errors on stability and performance of RC based control using force feedback. Our results indicate that a 1^st^ order RC improves the performance of force display, within the ranges of input period error of interest, more so than high-order RCs. Under nominal simulation and implementation conditions, the 1^st^ order RC option successfully reduced the 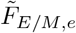 in the least number of cycles and to the lowest values.

We have found that the 3^rd^ order RC designed for random period error (RPE) only performed the best of all three RCs at higher half-width values in the random signal period error simulation analysis and not in the smaller half-width range examined in implementation. This findings is consistent with the sensitivity to random period error robustness analysis of various order RCs designed for random period error in previous work, in which a high-order RC with positive real fractional weights provides robustness to high values of RPE, also described as noise [28], [29]. We have found that 3^rd^ order RC designed for constant period error (CPE) only performed the best in the CPE simulation analysis in the region of mid-range frequency error values, in agreement with the CPE implementation results and sensitivity analysis (Fig. S4). These results are inconsistent with previous findings of 2^nd^ and 3^rd^ order RCs designed for CPE which reduce error at 0.5% constant period time error [22], a relatively small CPE value, and with previous sensitivity analyses [30]. However, the fact that the 3^rd^ order RC designed for CPE does not perform better at the smaller frequency error range, may simply be a result of the small magnitude of the selected RC gain and weights based on achievable stability and convergence in implementation.

Walking gait has a stride-by-stride variation in spatiotemporal parameters, or RPE, of 3-5% [20]. At the most comparable half-width of 0.04 Hz (8%) there is not a significant difference between the three RC types which all perform better than PPC. If the gait spatiotemporal parameter variation is considered in the context of constant offset from the fundamental frequency of the RC; at 0.02 Hz (4%) of frequency error, the RC-1 performs the best.

In previous lower extremity exoskeleton work, the quasi-periodic nature of gait has been exploited to reduce torque tracking error. Most notably, in a Bowden cable actuated ankle exoskeleton, the addition of iterative learning of desired motor position to proportional control with damping injection lead to stride-wise torque error reductions ranging between 38% and 84% [12]. Similarly, in the control of a lower extremity exoskeleton which added compensation for dynamics and feedforward torque tracking improved torque error at the hip and knee joints by at least 52% and 61%, respectively [31]. By comparison, under nominal conditions our 1^st^ order RC reduced 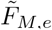 by 87%. Both under a half-width frequency condition of 0.04 Hz (8%) and at a constant frequency error of 0.02 Hz (4%), our 1^st^ order RC would still achieve a modeled reduction of 82%. It is important to note that our experimental platform is designed for upper extremity interaction, for which previous controllers have more varied results. In the pronation/supination joint of a wrist robot, the addition of gravity, inertia, and friction compensatory action on proportional control versus zero impedance PD control reduced the mean interaction torque by 77% [32]. More difficult to compare is the application of impedance compensation to an admittance controller of a hydraulic upper extremity exoskeleton; leading to a reduction of energy expenditure by the operator of up to 20% [33]. However, these two examples of upper extremity robotic controllers do not exploit the quasi-periodic nature of human performed tasks.

Considering the relatively small RPE and CPE ranges of interest given the spatiotemporal parameter variance of gait; we believe that 1^st^ order RC is the optimal RC design to apply in future work. Such work in the future will involve a robotic lower extremity exoskeleton platform designed for intervention utilizing existing force-feedback control [34].

---

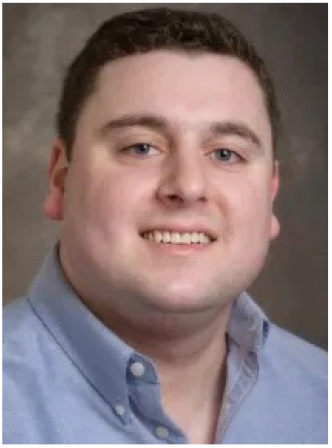

**Robert L. McGrath** (S’ 18) received the bachelor of mechanical engineering degree from the University of Delaware, Newark, DE, USA, in 2013. He is currently working towards the Ph.D. degree in biomedical engineering at the University of Delaware as a member of the Human Robotics Laboratory. His research interests include robotic lower extremity gait retraining, biomechanics, and repetitive control.

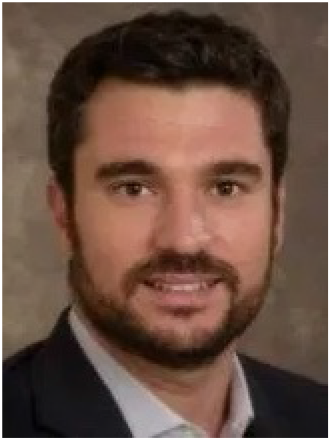

**Fabrizio Sergi** (M’ 13) received the B.S., M.S., and Ph.D. degrees in biomedical engineering from Universit’a Campus Bio-Medico di Roma, Rome, Italy, in 2005, 2007, and 2011, respectively. From 2012 to 2015, he was a postdoctoral fellow and research scientist at Rice University, Houston, Texas, where he conducted research in rehabilitation robotics. He is currently an Assistant Professor of Biomedical Engineering at the University of Delaware, where he directs the Human Robotics Laboratory. His research focuses on robotics for neurorehabilitation and human augmentation, haptics, and neuroimaging.

## V. Supplementary Materials

### A. Parameter Estimation

**Fig. S1:**
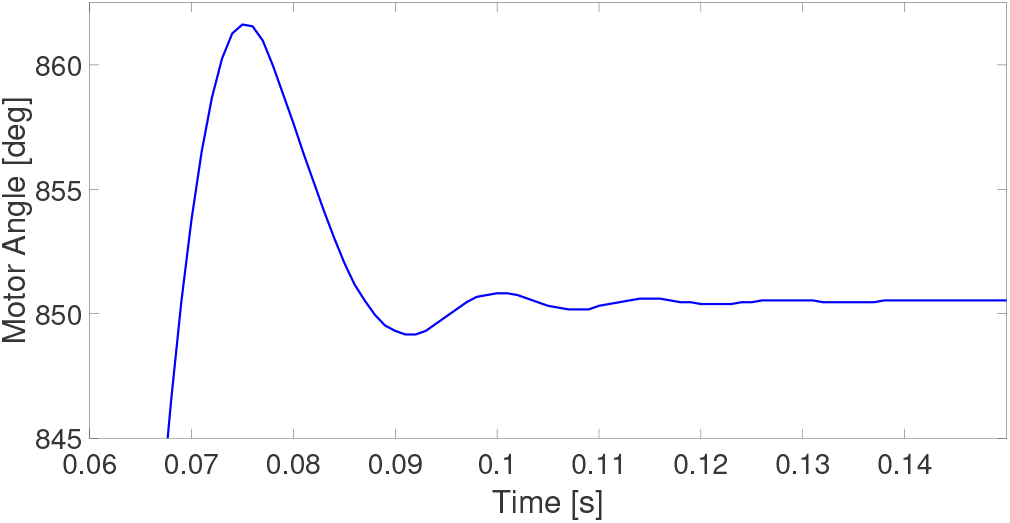
The open loop step response of the motor rotor with the end-effector in a blocked configuration. This motor angle response was utilized to estimate the natural frequency and damping ratio to derive the parameter estimates.

### B. Characterization

**Fig. S2:**
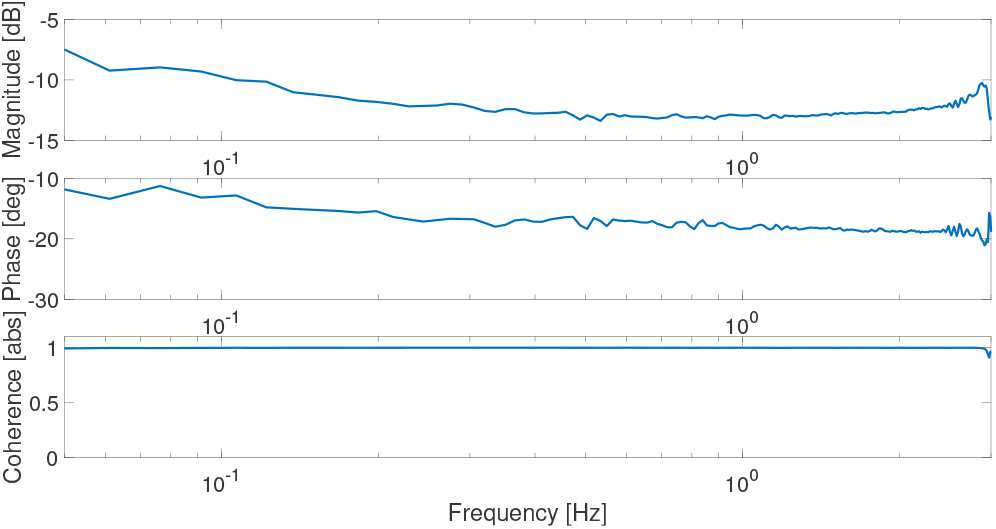
The magnitude, phase, and coherence of the characterized closed-loop transfer function of the robotic platform.

### C. FIR Filters

**Fig. S3:**
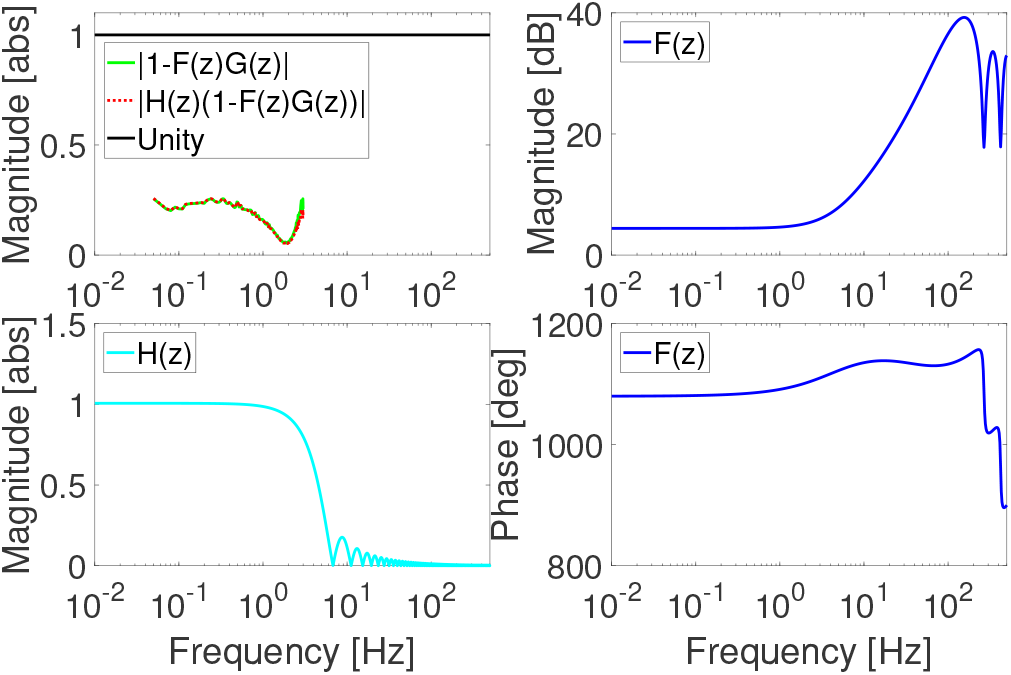
(Top left) Asymptotic stability of the compensator when coupled with the characterized plant and zero-phase low pass filter, (bottom left) the magnitude of the low pass filter, and (top right) the magnitude and (bottom right) phase of the compensator.

### D. Sensitivity

In Fig. S4 are the sensitivity magnitude plots of the four controllers. The plot describe the influence of the controller design on disturbance signals, such that a lower sensitivity, particularly in close proximity to the nominal frequency, is desirable.

**Fig. S4:**
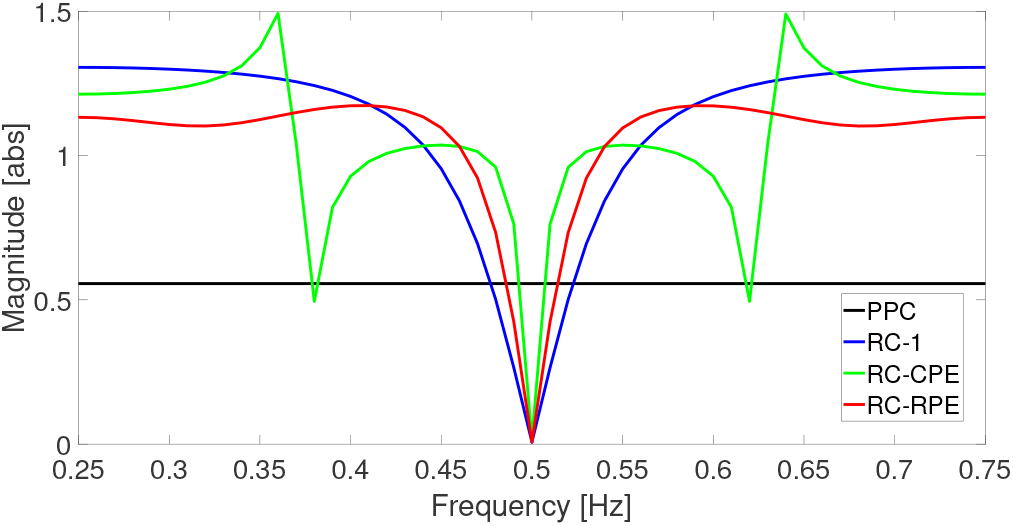
Magnitude plots for the sensitivity transfer functions of the four controllers.

